# Sleep Health in Lennox-Gastaut Syndrome impacts both Individuals with LGS and their Caregivers

**DOI:** 10.64898/2026.09.16.751819

**Authors:** Tracy Dixon-Salazar, Kathy J. Leavens, Helen Chen, Timothy J. Holick, Jeffrey A. Sweeney

## Abstract

Sleep disturbances are a significant but underrecognized burden for individuals with Lennox-Gastaut Syndrome (LGS) and their caregivers. We conducted a large LGS caregiver survey to assess sleep quality, duration, specific sleep-related issues, and management strategies. Quantitative analysis revealed a significant association between nocturnal seizures and overall seizure frequency, but general sleep metrics in individuals with LGS (e.g., sleep hours, sleep quality) were not associated with seizure burden. In contrast, caregiver sleep was frequently disrupted - particularly in families who co-slept or reported frequent seizures. Nearly 40% of caregivers experienced nightly sleep disturbances. Respondents reported a range of strategies to manage sleep problems, including medication, environmental adjustments, and behavioral routines. Many also expressed frustrations over the limited effectiveness of strategies and inadequate support for sleep issues. These findings underscore the dual impact of LGS-related sleep disturbances on patients and caregivers, and highlight the need for comprehensive, family-centered approaches to sleep health.

## Introduction

Lennox-Gastaut Syndrome (LGS) is a rare developmental and epileptic encephalopathy (DEE).^1,2^ LGS is diagnosed by two features observed via an electroencephalogram (EEG) recording, a slow spike-and-wave (SSW) signal and generalized paroxysmal fast activity (GPFA).^3,4^ Many patients with genetic DEE progress to LGS.^5–8^ LGS onset is typically before 5 years of age; LGS individuals experience multiple types of drug-resistant seizures, as well as impaired cognitive, behavioral, and communication development.^2^ LGS caregivers also experience significant physical and mental health burdens.^9–12^

Sleep disturbance and epilepsy have a bidirectional relationship.^13,14^ The frequency and type of seizures are influenced by the phase of sleep cycles. And both ictal and interictal epileptiform discharges interrupt sleep, and anti-seizure medications can alter sleep structure independent of seizures.^15,16^ In individuals with LGS, sleep disturbances are prevalent and often compound clinical challenges. The underlying epileptiform activity characteristic of LGS tends to intensify during certain stages of sleep, especially during non-rapid eye movement (NREM) phases, resulting in frequent arousals and fragmented sleep architecture.^17,18^ This disruption can exacerbate cognitive and behavioral difficulties, and reduce overall quality of life. Moreover, the interplay between nocturnal seizures, interictal discharges, and medication creates a complex landscape where restorative sleep is rarely achieved,^13,19^ underscoring the importance of comprehensive sleep assessment and management in LGS individuals.

We sought a caregiver perspective to better understand how seizures and sleep interact in LGS families. To advance patient-centered research in LGS, we prepared and distributed an online survey that queried seizure and sleep attributes of the LGS individual and the sleep of their caregivers.

## Results

### Survey respondents

A 25-question survey (Supplementary File 1) was designed to acquire information regarding sleep and seizure quality in LGS individuals and sleep quality in caregivers. There were 731 responses, of which 615 were complete and used for subsequent analyses. Respondents were directed to the survey primarily via advertising (Google search, 28%; Facebook, 21%) or the LGS Foundation website (www.lgsfoundation.org, 24%) (Figure 1A). The majority (78%) of respondents were parents of LGS individuals (Figure 1B). Data regarding age, seizure frequency and sleep hours were collected in ordinal bins. Age and sleep hours responses were normally distributed across bins (Figure 1C, E); however, 53% (325/615) of responses indicated >20 seizures per month (Figure 1D).

**Figure 1.**
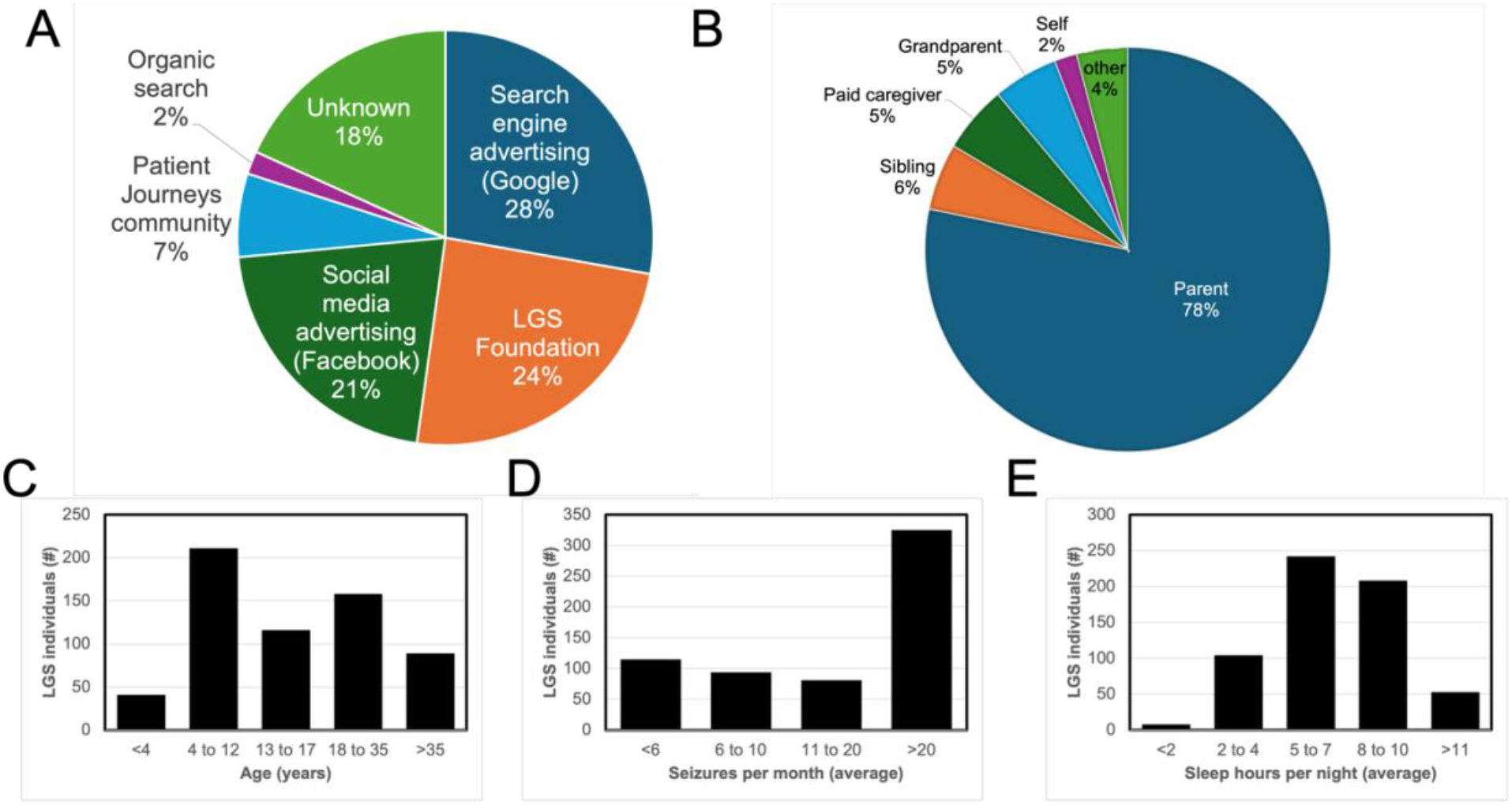
Summary of Survey Responses. **(A)** Respondents found the survey through diverse sources. **(B)** The majority of respondents were parents. **(C)** A broad age range of LGS individuals was surveyed. **(D)** Most of the LGS individuals surveyed have more than 20 seizures per month. **(E)** LGS individuals surveyed experience a broad range of sleep hours per night.

### Seizure frequency and sleep issues

In addition to general measures (e.g., age, seizure frequency, sleep hours per night), respondents were asked if specific sleep issues (falling asleep, staying asleep, nightmares or night terrors, nocturnal seizures, restless legs, sleep apnea, or other issues) were experienced by the LGS individual. We modeled sleep issues data using an ordinal logistic regression to determine if seizure frequency was predicative of specific sleep issues while testing age, sleep quality, and co-sleeping as covariates (Figure 2).

**Figure 2.**
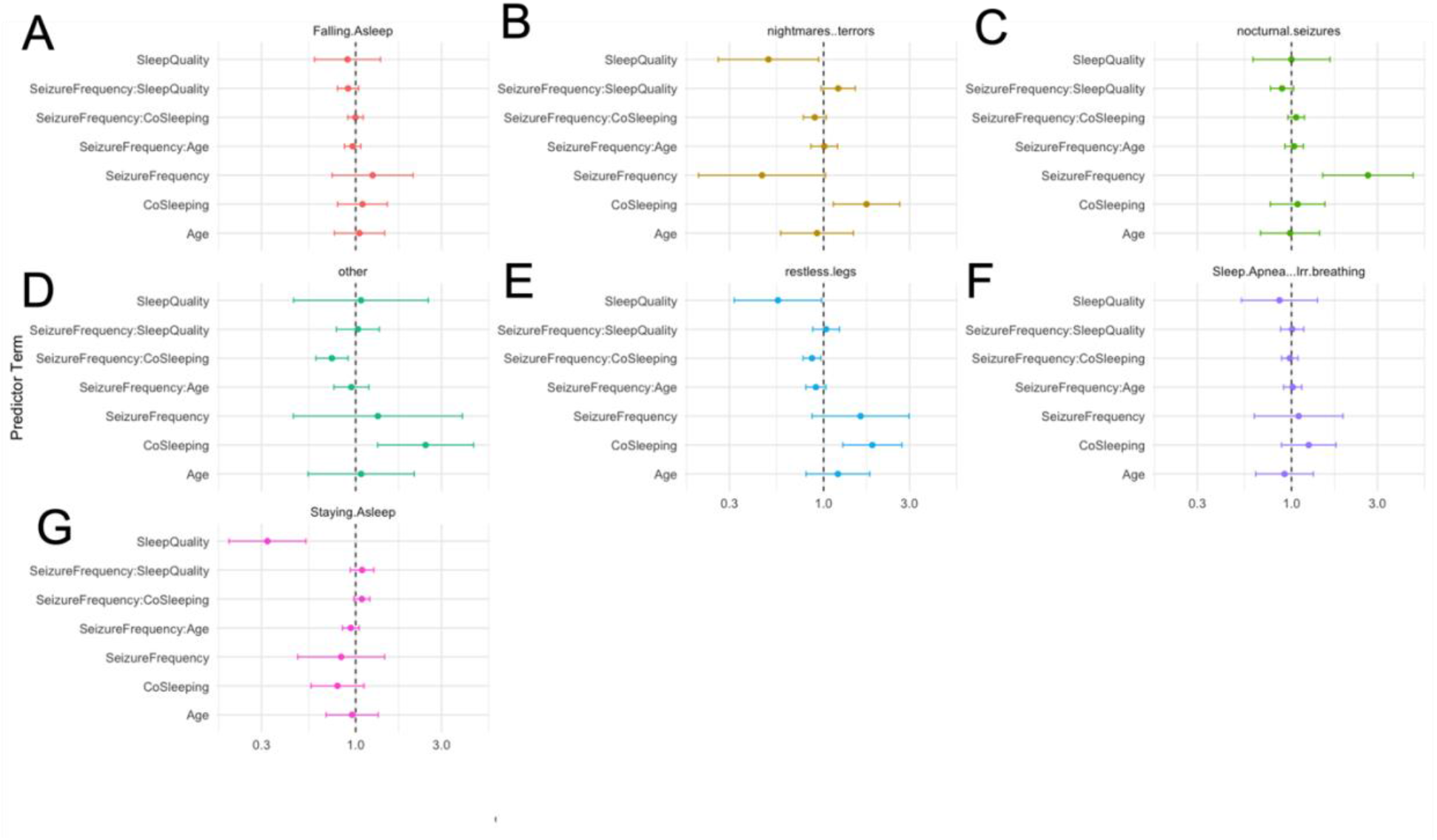
Association between individual LGS phenotypes and sleep-related Issues. **(A-G)** Ordinal logistic regression was performed considering seizure frequency, age, sleep quality, and co-sleeping as covariates; interactions between seizure frequency and age, sleep quality, and co-sleeping were also assessed (Y axis). The vertical, black, dotted line indicates the null hypothesis, statistical significance (p<0.05) is indicated when horizontal error bars are shifted left (negative association) or right (positive association) of null. (G) An expected strong negative association between sleep quality and staying asleep adds veracity to the overall approach. (B, D, E) Likewise, night terrors, restless legs, and the “other” phenotypes associate with co-sleeping given some likelihood of higher resolution observation in these individuals. (B, C) Seizure frequency is negatively associated with nighttime terrors and positively associated with nocturnal seizures.

Consistent with the expectation that seizures would impact sleep, a positive predictive association between seizure frequency and nocturnal seizures was observed (Figure 2C). The veracity of the approach is shown by the strong negative association between sleep quality and staying asleep (Figure 2G). Also, additional strong negative associations were observed between sleep quality and nightmares or night terrors (Figure 2B) and restless legs (Figure 2E). Co-sleeping showed strong positive associations with nightmares or night terrors (Figure 2B), restless legs (Figure 2E), and “other issues” (Figure 2D) possibly indicating better observation of some sleep issues by a co-sleeping caregiver.

### Caregiver reporting of seizures and sleep

We then sought to identify direct associations between caregiver reports of seizure and sleep measures. Seizure reports included measures of average seizure frequency per month (<6, 6 to 10, 11 to 20, and >20) and seizure control (1 (worst) to 5 (best)). Sleep reports likewise queried sleep in average hours per night (<2, 2 to 4, 5 to 7, 8 to 10, and >11) and sleep quality (1 (poor) to 5 (excellent)). Both measures of seizures and sleep correlated reasonably well with each other (Figure 3A, B).

**Figure 3.**
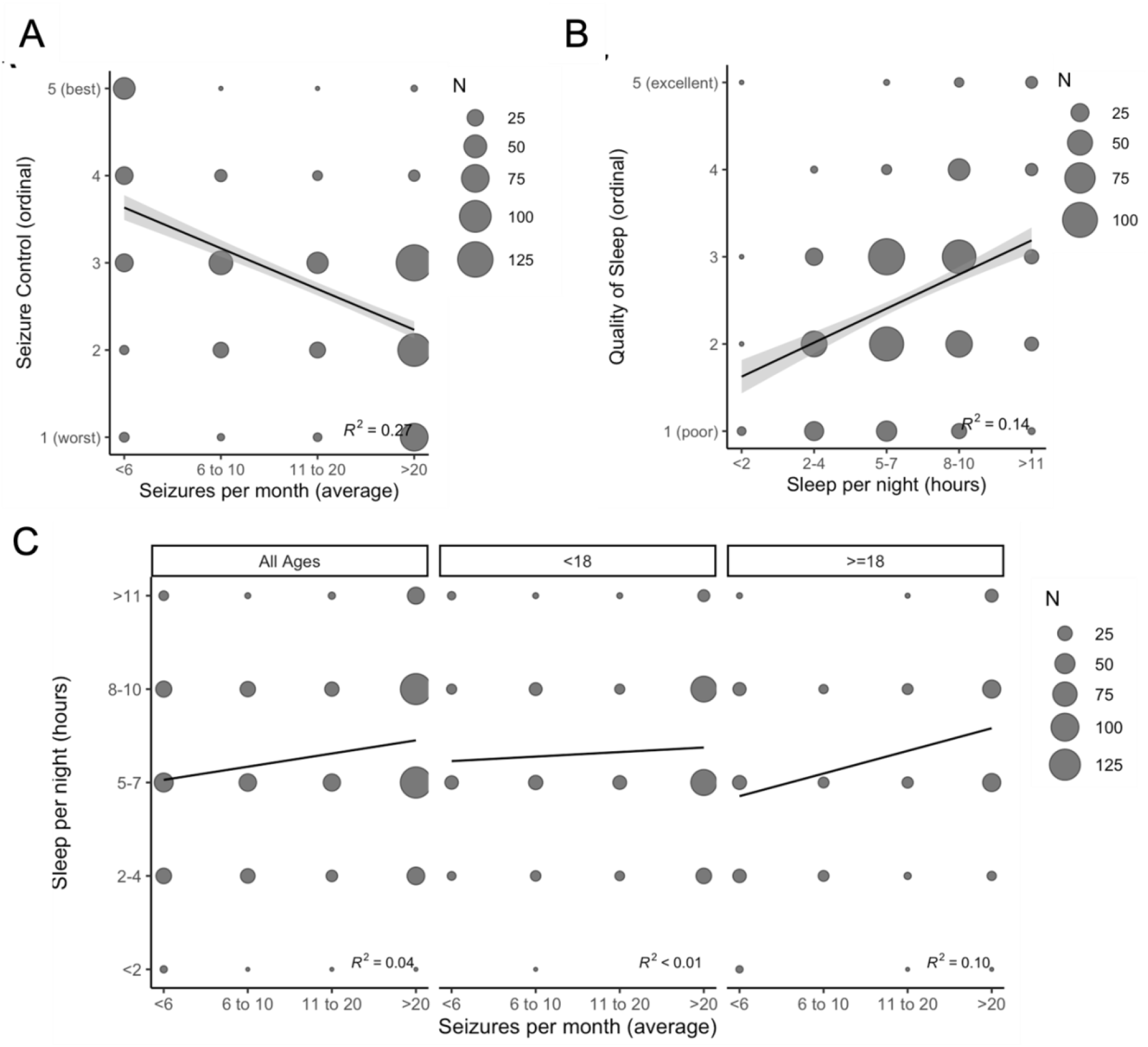
Respondent reports of seizures and sleep in LGS individuals. In all plots, the size of the marker indicates the number (N) of respondents. **(A)** Seizure control reported as an ordinal (1 (worst) to 5 (best), y axis) correlates weakly (R2 = 0.27) with reported frequency of seizures per month (x axis). **(B)** Quality of sleep reported as an ordinal (1 (poor) to 5 (excellent), y axis) also correlates weakly (R2 = 0.14) with reported sleep hours per night (x axis). **(C)** No significant correlation between sleep hours (y axis) and seizure frequency (x axis) was observed (left panel), even when age (<>18) is considered (center and right panels).

Interestingly, caregiver reporting of seizure frequency and sleep hours did not show expected significant associations (R^2^ = 0.04), even when age groups (<18, =>18) were analyzed separately (Figure 3C). Future survey designs targeting the LGS population should be designed with finer grained resolution of seizure frequency (e.g., seizures per week rather than seizures per month) to avoid dense clustering of responses in the highest (>20 seizures per month) frequency group (Figure 1D), as the majority of LGS patients experience >20 seizures per month with minimal seizure control. This is also illustrated in Figure 3A where individuals experiencing >20 seizures per month had seizure control reported as a 3, 4, or 5.

### Caregiver sleep and co-sleeping

The survey also asked respondents about their sleep, 84% (515/615) responded to this query. The majority (78%) of respondents indicated that their sleep was disturbed at least 1-2 times per week with 40% (204/515) indicating that their sleep is disturbed every night. Furthermore, we find a significant negative effect of co-sleeping on caregiver sleep (Figure 4A, p<0.0001). Co-sleeping was also significantly associated with increased seizure frequency (Figure 4A, p = 0.032) and mildly associated with fewer sleep hours for the LGS individual (Figure 4A, p = 0.084). As noted regarding an association between co-sleeping and specific sleep disturbances (Figure 2B, D, and E), co-sleeping caregivers may have better awareness of night time seizures and sleep hours in LGS individuals.

**Figure 4.**
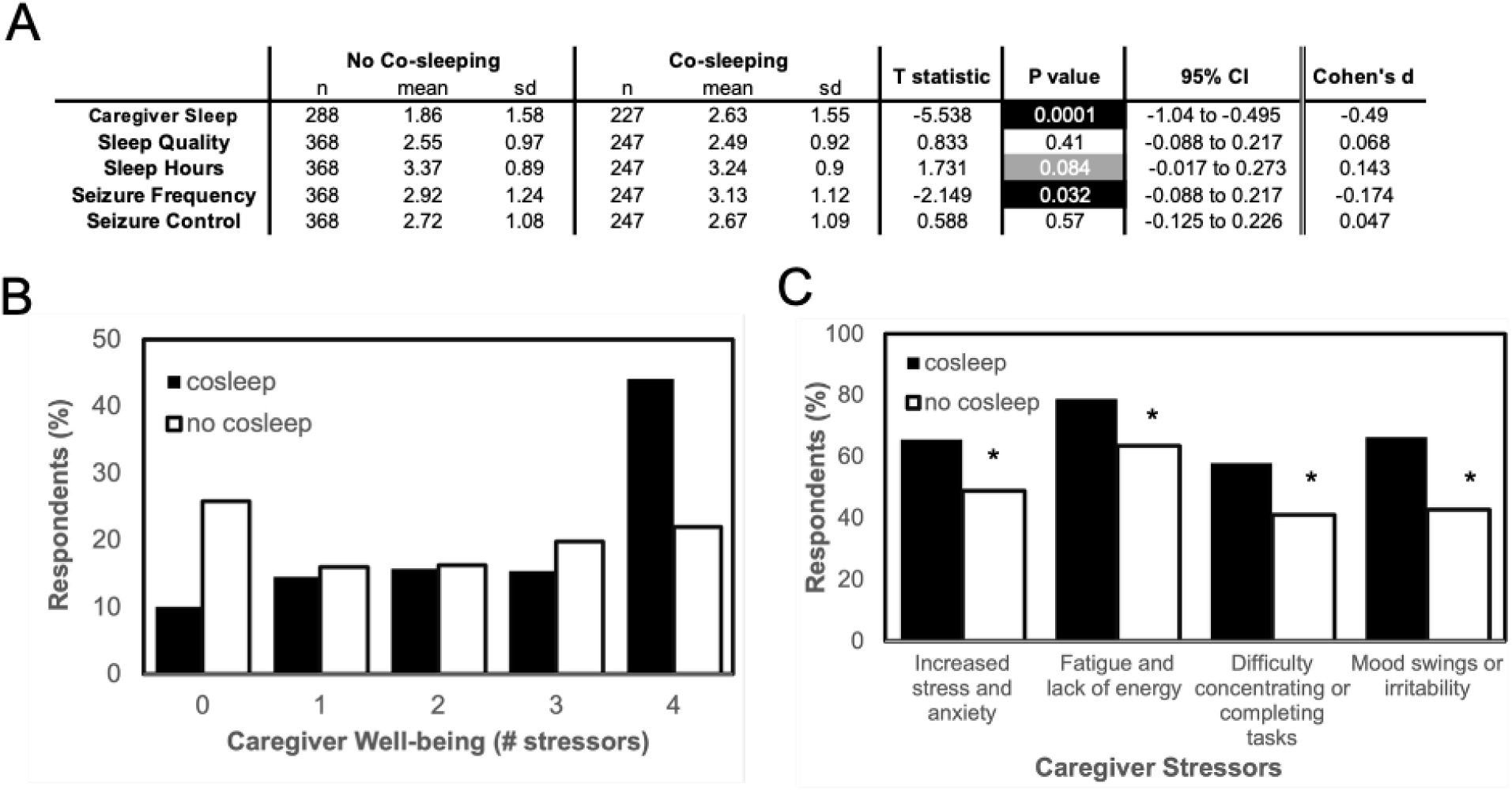
Impact of Co-sleeping on Caregiver Sleep and Well-being in LGS. **(A)** Group-level comparisons of caregiver-reported sleep, seizure outcomes, and seizure control in families who do not co-sleep versus those who do co-sleep. Co-sleeping was associated with significantly less caregiver sleep (p < 0.0001) and higher reported seizure frequency (p = 0.032). Data are presented as mean ± standard deviation (SD), with T statistics, p-values, confidence intervals (CI), and Cohen’s d effect sizes shown for each comparison. **(B)** Distribution of caregiver well-being based on the number of stressors reported. Caregivers who co-slept reported higher numbers of stressors. **(C)** Percentage of respondents reporting individual caregiver stressors. Co-sleeping caregivers were significantly more likely to report increased levels of all stressors (*, X^2^ p < 0.0001).

Caregivers were asked about sleep-associated factors that affected their well-being, such as increased stress and anxiety, fatigue and lack of energy, difficulty concentrating or completing tasks, and mood swings or irritability. Co-sleeping caregivers often reported that all four stressors affected their well-being (Figure 4B). Each individual stressor was reported significantly (X^2^, p value <0.0001) more by caregivers that co-slept (Figure 4C).

### Sleep management

The survey also asked about factors that worsen sleep-related issues and the approaches employed by caregivers to address sleep issues in LGS individuals (Figure 5). Implementing sleep hygiene practices and making medication adjustments are the methods most often used to address sleep-related issues. While the methods used to address sleep-related issues are similar by age, use of video camera monitoring and assistive devices is reported more frequently for patients under 18. Over half of respondents report they have experienced challenges or difficulties while trying to implement strategies to address sleep-related issues for the LGS patient. Belief that managing seizures is the key for better sleep, cost and financial barriers, and perception that there are no new solutions that would address sleep-related issues dissuade caregivers from seeking professional help.

**Figure 5.**
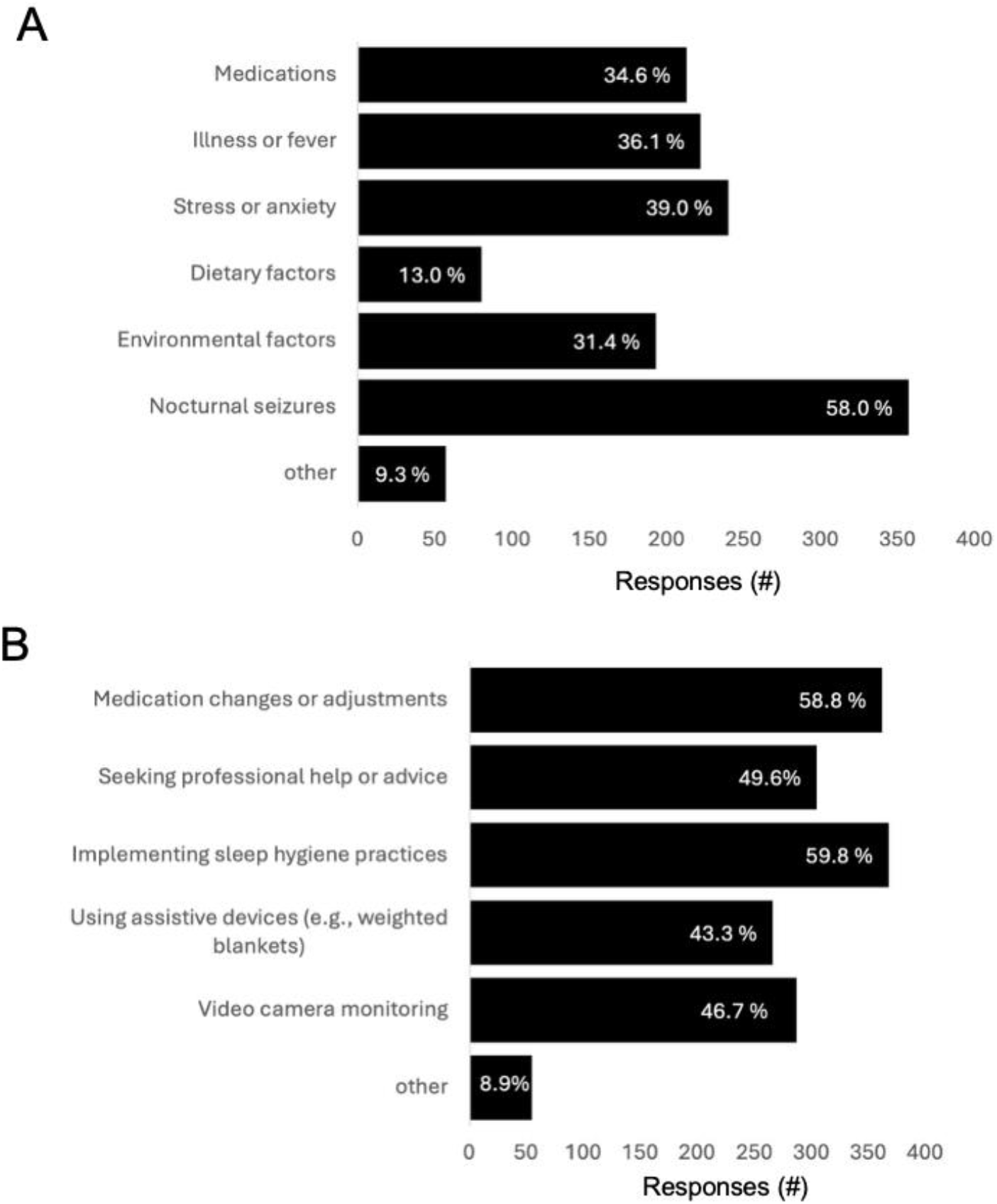
Qualitative factors reported by caregivers. **(A)** Factors that are observed to worsen sleep-related issues in LGS individuals. Factors are listed on the y-axis, bar length indicates the number of respondents (x-axis) reporting that the factor worsens sleep-related issues. The percent of total responses (n = 615) is indicated on each bar. **(B)** Approaches used to address sleep-related issues. Approaches are listed on the y-axis, bar length indicates the number of respondents (x-axis) that have employed each approach. The percent of total responses (n = 615) is indicated on each bar.

## Discussion

Caregiver survey data can provide important insights for patient-centered outcomes research in LGS.^20,21^ We found that specific sleep-related issues, nocturnal seizures, nighttime terrors, and restless legs, were associated with increased seizure frequency. Perhaps not unexpectedly, co-sleeping diminished the quantity of caregiver sleep. Co-sleeping also contributed to the reporting of significant sleep-related issues, and co-sleeping caregivers reported associations between fewer seizures and better sleep that were not apparent when all caregiver reported data was analyzed.

Many respondents mentioned seeking guidance from neurologists, sleep specialists, or other medical professionals. The most frequently reported sleep advice involved adjusting medications to help manage sleep, including the use of melatonin, clonidine, or trazodone. Caregivers were often advised to try different doses or combinations of these medications to improve sleep. Caregivers were also advised to implement sleep hygiene practices, including establishing consistent bedtime routines, avoiding screens before bed, and creating a calm and relaxing environment before sleep. Several families were advised to pursue medical tests like sleep studies to identify underlying issues such as sleep apnea. CPAP machines and seizure monitors were also recommended as part of sleep management strategies. Some respondents expressed frustration, stating that despite seeking professional help, they found the advice ineffective in managing sleep issues. This was especially the case for those dealing with nocturnal seizures.

Many caregivers indicated that managing seizures is their top priority. They often feel that sleep issues are secondary and related to seizures, so they prefer to focus on seizure control rather than seeking separate sleep-specific help. The high cost of professional services, combined with inadequate insurance coverage, was a significant reason caregivers avoided seeking help. Some mentioned that the expense was simply too high for them to pursue additional assistance. Some respondents also expressed doubt that professional help would provide new or effective solutions, especially if they have already tried medical interventions that didn’t work. As a result, they chose not to seek further help. Several respondents were hesitant to seek professional advice because they did not want to add more medications, especially if they felt their loved ones were already on many drugs with concerning side effects. A portion of respondents were simply unaware that professional help for sleep issues was available or didn’t know which type of specialist to consult.

Respondents mentioned a combination of using medications, environmental changes, and behavioral adjustments as common approaches to improving sleep quality for caregivers. Many respondents mentioned using medications such as melatonin, over-the-counter sleep aids, or prescription medications like trazodone to help them sleep better. Several caregivers shared that maintaining a consistent bedtime routine, including relaxing activities like taking a warm bath, listening to soothing music, or dimming the lights, helped them improve their sleep quality. Physical activities like exercising during the day, yoga, or taking a walk before bed were mentioned by respondents as helpful strategies to tire the body and promote better sleep. Adjusting the sleep environment, such as using white noise machines, blackout curtains, or ensuring the room is cool and quiet, was frequently highlighted as a key factor in improving sleep. Some caregivers reported that taking naps during the day or alternating night duties with a partner helped them manage the challenges of disrupted sleep, ensuring they got rest when possible.

Moving forward, this study adds to the growing body of research on caregiver-reported outcomes in LGS. By capturing caregivers’ experiences and strategies related to sleep and seizure management, our findings highlight both the complexity of caregiving and the need for integrated clinical guidance that addresses sleep alongside seizure control. Future research and clinical interventions should consider the bidirectional relationship between sleep and seizures, the unique challenges of co-sleeping, and barriers to accessing professional help. Tailored support that combines medical, behavioral, and environmental strategies will be beneficial to the well-being of both LGS individuals and their caregivers.

## Acknowledgements

The authors would like to thank the patients and their families who participated in this initiative, as well as the staff of the LGS Foundation and PatientJourneys.org for their support. Manuscript preparation and data analysis was provided by Michael J. McConnell, PhD (Rare Mosaic scientific consulting, Charlottesville, VA). Additional data analysis support was provided by Juan Falla (Nutley, NJ) and Jessica Maginski (JM Research Solutions, Inc., Charleston, SC). The authors would also like to thank Jazz Pharmaceuticals, Inc. for funding this initiative.

## Methods

### Survey Design

The LGS Sleep Health Survey was developed by Patient Journeys in collaboration with the LGS Foundation. Survey questions were informed by input from LGS caregivers and healthcare professionals and were guided by industry standards for patient-focused data collection.

The survey was available online from December 15, 2023, through May 9, 2024 (160 days). Data was collected anonymously. While participants who completed the survey and provided an email address received a $25 honorarium, compensation was not mentioned in any advertising materials to preserve data integrity. Email addresses were not linked to individual survey responses.

### Data Analysis

Survey data was consolidated and initially analyzed using Excel (Microsoft, Redman, WA). Subsequent analyses were performed in R^22^ with code suggestions generated using prompts in ChatGPTv4 (Open AI) and Claude (Anthropic). Ordinal logistic regression was performed using the MASS^23^ and tidyverse^24^ packages. Figure panels were prepared using Excel and ggplot, and assembled in Powerpoint (Microsoft).

