## Supplemental File 1 for "Sleep Health in Lennox-Gastaut Syndrome impacts both Individuals with LGS and their Caregivers"

### LGS Sleep Survey

**1. What is your relationship to the person with LGS?**

- [ ] Parent
- [ ] Grandparent
- [ ] Sibling
- [ ] Paid caregiver
- [ ] Other

**2. How old is the person with LGS?**

- [ ] Under 4
- [ ] 4-12
- [ ] 13-17
- [ ] 18-35
- [ ] Over 35

**3. Before we ask about sleep-related matters, we have a few general questions regarding seizures. How effectively are y our loved one's seizures currently being controlled?**

| 1 | 2 | 3 | 4 | 5 |
|---|---|---|---|---|
| [ ] | [ ] | [ ] | [ ] | [ ] |

Not at all Moderately Very well

**4. How many seizures does your loved one experience in an average month?**

- [ ] 5 or less
- [ ] 6-10
- [ ] 11-20
- [ ] More than 20

**5. Now, let's move on to our sleep-related questions. On average, how many hours of sleep per night does your loved on e typically get?**

- [ ] Less than 2
- [ ] 2-4
- [ ] 5-7
- [ ] 8-10
- [ ] 11 or more

**6. How would you rate their overall quality of their sleep, (did they get a good night's sleep)?**

| 1 | 2 | 3 | 4 | 5 |
|---|---|---|---|---|
| ☐ | ☐ | ☐ | ☐ | ☐ |
| Poor | | Average | | Excellent |

**7. What specific sleep-related issues, if any, does your loved one experience?**

- [ ] Difficulty falling asleep
- [ ] Difficulty staying asleep
- [ ] Sleep apnea or irregular breathing
- [ ] Nightmares or night terrors
- [ ] Restless legs or periodic limb movements
- [ ] Sleepwalking
- [ ] Nocturnal seizures
- [ ] Bed wetting
- [ ] Daytime napping
- [ ] Other

**8. In an average week, how many times does your loved one have dificulty falling asleep?**

- [ ] 1-2
- [ ] 3-4
- [ ] 5 or more

**9. In an average week, how many times does your loved one have dificulty staying asleep?**

- [ ] 1-2
- [ ] 3-4
- [ ] 5 or more

**10. In an average week, how many times does your loved one experience Sleep apnea or irregular breathing?**

- [ ] 1-2
- [ ] 3-4
- [ ] 5 or more

**11. In an average week, how many times does your loved one experience nightmares or night terrors?**

- [ ] 1-2
- [ ] 3-4
- [ ] 5 or more

**12. In an average week, how many times does your loved one experience restless legs or periodic limb movements?**

- [ ] 1-2
- [ ] 3-4
- [ ] 5 or more

**13. In an average week, how many times does your loved one experience sleepwalking?**

- [ ] 1-2
- [ ] 3-4
- [ ] 5 or more

**14. In an average week, how many times does your loved one experience nocturnal seizures?**

- [ ] 1-2
- [ ] 3-4
- [ ] 5 or more

**15. In an average week, how many times does your loved one experience bed wetting?**

- [ ] 1-2
- [ ] 3-4
- [ ] 5 or more

**16. In an average week, how many times does your loved one experience daytime napping?**

- [ ] 1-2
- [ ] 3-4
- [ ] 5 or more

**17. In an average week, how many times does your loved one experience these other sleep-related issues?**

- [ ] 1-2
- [ ] 3-4
- [ ] 5 or more

**18. Which of the factors listed below, if any, do you believe worsen your loved one's sleep-related issues?**

- [ ] Medications
- [ ] Illness or fever
- [ ] Stress or anxiety
- [ ] Dietary factors
- [ ] Environmental factors (noise, light, temperature)
- [ ] Nocturnal seizures
- [ ] Other

**19. Do you, or any other caregivers in the home, co-sleep with the person who has LGS?**

- [ ] Yes
- [ ] No

**20. In an average week, how many times does someone co-sleep with the person?**

- [ ] 1-2
- [ ] 3-4
- [ ] 5 or more

**21. Which of these things, if any, have you tried to address the sleep-related issues?**

- [ ] Medication changes or adjustments
- [ ] Seeking professional help or advice
- [ ] Implementing sleep hygiene practices, (e.g., consistent schedule, adjusting before bedtime behaviors, relaxation wind down periods)
- [ ] Using assistive devices (e.g., weighted blankets)
- [ ] Video camera monitoring
- [ ] Other

**22. What professional help or advice did you receive?**

**23. Is there any particular reason you have not sought professional help to address sleep issues?**

**24. Have you encountered any challenges or difficulties while implementing strategies to address sleep-related issues?**

- [ ] Yes
- [ ] No

**25. Would you describe the challenges or difficulties you faced?**

**26. Are there any other interventions or strategies that you are considering to address your loved one's sleep issues?**

**27. Do you have any further comments or insights you would like to share about ideas used to address sleep issues in in dividuals with LGS?**

**28. We're almost finished! Now we'd like to inquire about the quality of your sleep as a caregiver. During an average wee k, how many nights do you encounter sleep disruptions because of your loved one's seizure disorder?**

- [ ] 0 (my sleep is not disturbed)
- [ ] 1-2
- [ ] 3-4
- [ ] 5-6
- [ ] Every night

**29. How do sleep disruptions impact your overall well-being?**

- [ ] Increased stress and anxiety
- [ ] Fatigue and lack of energy
- [ ] Difficulty concentrating or completing tasks
- [ ] Mood swings or irritability
- [ ] Other

**30. What things do you do to help you get a better night's sleep?**

**31. Thank you for completing this survey on the Impact of Sleep on Individuals with Lennox-Gastaut Syndrome! Please provide your email below, and we will send you the $25 Amazon gift card!**

(By providing your email, you agree to our Terms and Conditions and consent to our Privacy Policy.)
